# Lysosomal MLKL is balanced by ESCRT to control cell death

**DOI:** 10.1101/2023.08.29.555049

**Authors:** Catherine Jamard, Clara Gil, Marie Castets, Gabriel Ichim, Kathrin Weber

## Abstract

Mixed lineage kinase-like (MLKL) is activated by RHIM-domain containing kinase (RIPK)3 to permeabilize the plasma-membrane and execute necroptosis, a form of regulated necrosis. We found that MLKL is activated in an atypical, RIPK3- and necroptosis-independent manner downstream of Toll-like receptor 3, resulting in its translocation to lysosomes and lysosomal membrane permeabilization. Damaged lysosomes then undergo exocytosis, leading to the integration of lysosomal MLKL into the plasma-membrane to trigger cell death. The ESCRT-machinery can repair damaged lysosomes and counteract cell death by packing lysosomal MLKL into intraluminal vesicles, which are subsequently released as extracellular vesicles. In this way, ESCRT-machinery balances life and death decisions by preventing lysosomal MLKL to reach its killing destination, which is the plasma-membrane.

## Introduction

Beyond apoptosis, several forms of necrotic cell death are also regulated by genetically encoded signaling pathways ^1^. Necroptosis is one of the best-characterized forms of regulated necrosis, and relies on the plasma-membrane permeabilizing activity of mixed lineage kinase domain-like pseudokinase (MLKL). Following its phosphorylation by RHIM-domain containing kinase (RIPK)3 within its activation loop, a conformational change unleashes the N-terminal four-helical bundle domain (4HBD) from the C-terminal pseudokinase domain (PsKD) of MLKL. Oligomerization into different multimeric species and the translocation to the plasma-membrane occur subsequently before MLKL disrupts the plasma-membrane leading to cellular swelling and cell death ^2–5^.

During necroptosis MLKL-mediated plasma-membrane lesions can be repaired by the activity of the endosomal sorting complex required for transport (ESCRT) machinery ^6^. By shedding of plasma-membrane vesicles, which contain active MLKL, the ESCRT machinery delays necroptosis and prolongs cell survival. Next to the plasma-membrane, ESCRT was also shown to repair small lysosomal membrane ruptures, however, the underlying mechanism remains unanswered ^7,8^.

Although the killing function of MLKL is best characterized in TNFR-induced necroptosis, it can be also be activated by the innate immune receptor Toll-like receptor (TLR)3 ^9^. Recently, we could further expand the repertoire TLR3-induced cell death modalities to lysosomal cell death (TLR3-LCD), which occurred in the absence of Caspase-8 (apoptosis) and RIPK3 (necroptosis) in the neuroblastoma cell line SH-SY5Y ^10^. Although TLR3-LCD resembles classical features of LCD with lysosomal membrane permeabilization (LMP) being the initiating event, the exclusive role of cathepsins in cell death induction could not be demonstrated and the underlying molecular mechanism remains unclear.

## Results

### Lysosomal MLKL is required to execute TLR3-LCD

MLKL is an interferon-sensitive gene (ISG) ^11^. Type I IFN (IFN) and/or pIC treatment upregulated MLKL in SH-SY5Y cells in a time- and treatment dependent manner (Fig. 1a; Suppl. Figure S1a), pointing towards a potential role of MLKL in execution of TLR3/LCD. CRISPR/Cas9 mediated MLKL knock-out by two different guide RNAs significantly decreased IFN/pIC induced cell death indicating the crucial role of MLKL in TLR3-LCD execution (Fig. 1b and c). Further, the inhibitor of active MLKL, necrosulfonamide (NSA), reduced IFN/pIC-induced cell death (Fig. 1d). Importantly, inhibition of the necroptotic activator of MLKL, RIPK3 by GSK-872’, had no effect on TLR3-LCD induction, which is in line with the lack of RIPK3 expression in these cells (Fig. 1a and d). Besides necroptosis, NSA can also inhibit pyroptosis by binding to Gasdermin D ^12^. However, TLR3-induced pyroptosis requires Caspase-8 which is not expressed in SH-SY5Y cells ^10,13^. Thus, together these results point towards a RIPK3/necroptosis-independent role of MLKL in TLR3-LCD execution. To analyze MLKL activation during TLR3-LCD, we used Phostag SDS-PAGE. Distinct slower migrating MLKL species were detectable following IFN/pIC treatment, which were not present after lambda phosphatase treatment (Fig. 1e). Importantly, no signal was obtained when an antibody directed against the necroptotic MLKL phosphorylation site(s), T357/S358, was used. Thus, during TLR3-LCD MLKL is phosphorylated at necroptotic-distinct site(s) and by another kinase than RIPK3. Following IFN/pIC treatment, MLKL also assembled into higher-order complexes detectable as a smear on non-reducing SDS-PAGE (Fig. 1f). A proportion of MLKL oligomers appears to be too large to enter the stacking gel and accumulate at the bottom of the well. DTT collapsed the smear, indicating its specificity (Fig. 1f, left panel). This oligomeric pattern of MLKL is in sharp contrast to the necroptotic one ^2^ (Fig. 1f, right panel). To fulfill its membrane permeabilizing function, MLKL has to translocate to membranes and we found that MLKL re-located from a diffuse to a cytosolic, punctate pattern following IFN/pIC treatment (Fig. 1g). These MLKL puncta co-localized with the lysosomal marker protein LAMP-1 and the percentage of LAMP-1, which overlaid with MLKL increased significantly with IFN/pIC treatment (Fig. 1h). Thus, besides its atypical, RIPK3/necroptosis-independent activation, MLKL translocated to lysosomes during TLR3-LCD.

**Fig. 1.**
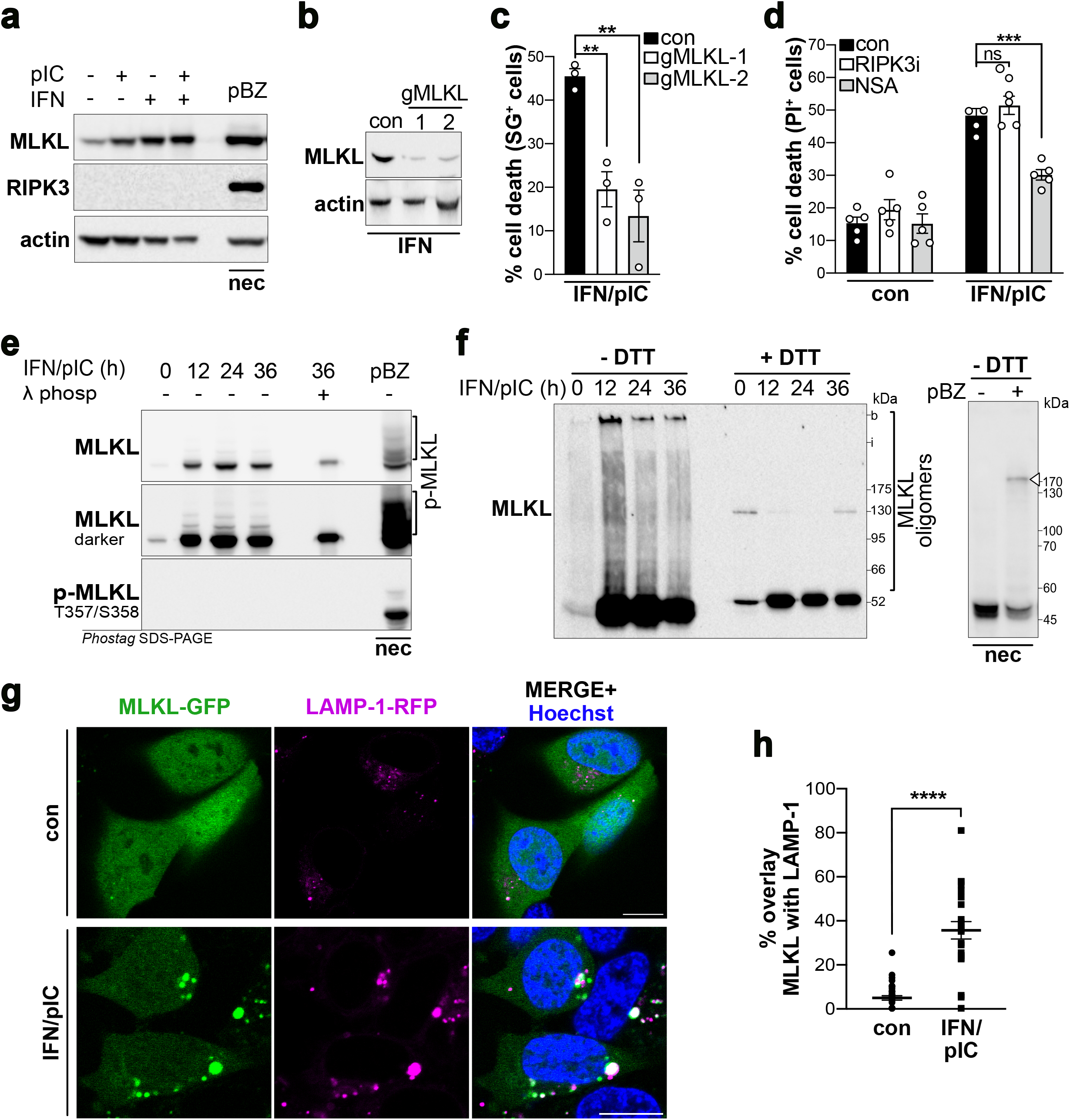
Lysosomal MLKL is required to execute TLR3-LCD. (**a**) Western blot analysis of SH-SY5Y cells primed for 16 h with IFN type I followed or not by treatment with Poly(I:C) (IFN/pIC) for 48 h using anti-MLKL, -RIPK3 and -actin antibodies. (**b** and **c**) MLKL was knocked out by CRISPR/Cas9 approach using two different gRNAs in SH-SY5Y cells. (**b**) Western blot analysis of MLKL expression levels using anti-MLKL and -actin antibodies treated with IFN for 16 h. (**c**) Cell death profile following IFN/pIC treatment for 48 h analyzing SytoxGreen (SG) positive cells. Data points represent the mean ± □S.E.M. of n = 3. **p = 0.009 and 0.0032. (**d**) Cell death profile analyzing PI positive SH-SY5Y cells, left untreated or treated for 48 h with IFN/pIC in combination with GSK-872’ (RIPK3i) or necrosulfonamide (NSA). Data points represent the mean ± S.E.M. of four independent experiments (n = 4); ***p=0.0004; ns= not significant. (**e**) Phostag SDS-Page of SH-SY5Y cells treated for indicated times with IFN/pIC or HT-29 cells with pBZ for 12 h to induce necroptosis using anti-p-MLKL (T357/S358) or anti-MLKL antibody. Phospho-MLKL bands were verified by lambda phosphatase treatment. A representative of n = 3 is shown. (**f**) Western blot analysis under non-reducing (-DTT) and reducing (+DTT) conditions of SH-SY5Y cells treated with IFN/pIC for the indicated times using an anti-MLKL antibody. A representative of n=3 is shown; left panel. Oligomeric MLKL is indicated. Right panel: Western blot analysis under non-reducing conditions of HT-29 cells treated with pBZ (poly(I:C), BV6 and zVAD-fmk; necroptosis (nec)) for 12 h using an anti-MLKL antibody. Oligomeric MLKL is indicated by an arrowhead. b: bottom of the well; i: interface between stacking and running gel. (**g**) Confocal images of MLKL-GFP expressing SH-SY5Y cells expressing LAMP-1-RFP and Hoechst left untreated or treated for 24 h with IFN/pIC. Merged image of MLKL-GFP, LAMP-1 and Hoechst right panel. Scale bars 10 µm. Representative images of n = 1 are shown. (**h**) Quantification of (**g**) analyzing the percentage of overlay of LAMP-1-RFP fluorescence intensity with MLKL-GFP. Data points represent the mean ± S.E.M. of n = 2 analyzing at least 25 cells. ****p < 0.0001.

### Lysosomal exocytosis enhances TLR3-LCD

The N-terminal four helical bundle domain (4HBD) of MLKL is attributed with its membrane permeabilizing function, which can be abolished by N-terminal addition of a strep-affinity-tag (strep-MLKL) or deletion of the 4HBD (PsKD) ^3,5^. Both dominant negative mutants failed to translocate to lysosomes following IFN/pIC treatment, which resulted in significant reduction of LMP as well as cell death (Fig. 2a; Suppl. Figure S2a). Thus, atypical, RIPK3/necroptosis-independent activated MLKL mediates LMP. However, cathepsins did not significantly contributed to TLR3-LCD execution, suggesting that MLKL-mediated LMP is limited and the release of lethal lysosomal content into the cytoplasm is prevented (Suppl. Figure S2b). Alternatively, cell death execution was previously associated with lysosomal exocytosis, which is the fusion of lysosomes with the plasma-membrane ^14^. In SH-SY5Y, like in most in non-polarized cells, lysosomes mainly concentrate in the perinuclear region (Fig. 2b and c), making their fusion with the plasma-membrane unlikely. In contrast, IFN/pIC treatment inverted this intracellular distribution and lysosomes re-distributed to the cell periphery. Of note, NSA prevented this re-distribution highlighting its dependency on active MLKL (Fig. 2c). We next tested whether peripheral lysosomes undergo lysosomal exocytosis. The appearance of integral lysosomal proteins at the plasma-membrane can be used as a read-out for lysosomal exocytosis. Surface staining of the lysosomal protein LAMP-1 revealed an overall increase of lysosomal exocytosis during TLR3-LCD (Fig. 2d and e). Immunoprecipitation of surface proteins confirmed the increase of LAMP-1 on the cell surface during TLR3-LCD (Fig. 2f). Besides LAMP-1, also MLKL was detectable in surface IPs following IFN/pIC treatment, which supports the idea that lysosomal MLKL appears at the plasma-membrane through lysosomal exocytosis (Fig. 2f). Further, we observed that following IFN/pIC treatment dying cells inflate (Suppl. Figure S3a and b). Considering that lysosomal MLKL exerts membrane permeabilizing capacities, its integration into the plasma-membrane might induce cellular swelling and explosion and finally cell death ^15^. To pinpoint the pro-death role of lysosomal exocytosis during TLR3-LCD, we used MK6-83, an accelerator of lysosomal movement to the cell periphery and lysosomal exocytosis (Suppl. Figure S4a, b and c) ^16,17^. MK6-83 significantly increased IFN/pIC induced cell death (Fig. 2g). Conversely, we used the two known inhibitors of lysosomal exocytosis LY294002 and Vacuolin-1 ^18,19^. While Vacuolin-1 treatment resulted in the appearance of previously described LAMP-1-positive vacuoles, treatment with LY294002 resulted in significant increase in central clustering of lysosomes (Suppl. Figure S4d and e). Both inhibitors significantly reduced IFN/pIC induced cell death (Fig. 2h), reinforcing the idea that lysosomal MLKL exerts its deadly role at the plasma-membrane.

**Fig. 2.**
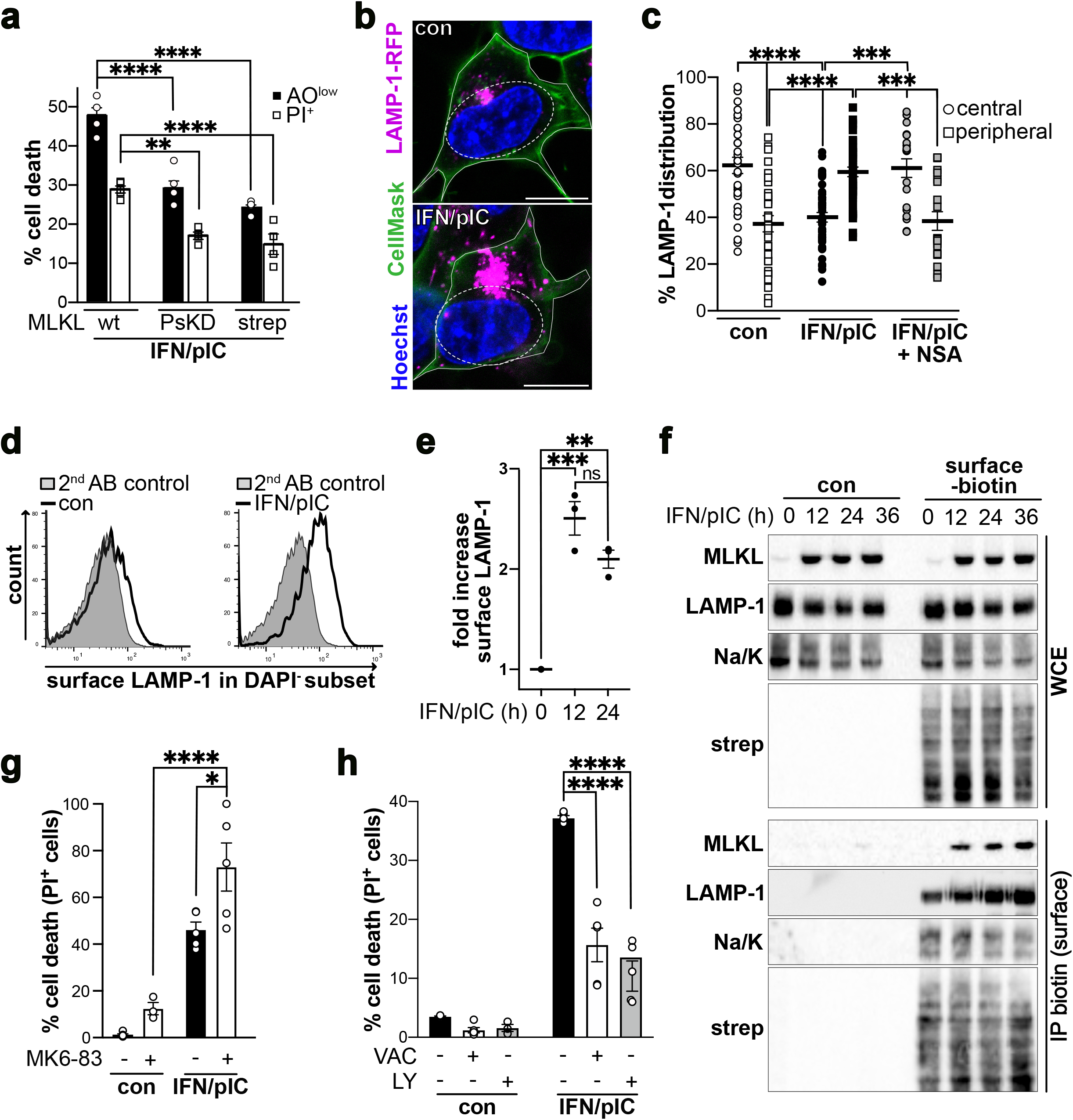
Lysosomal exocytosis enhances TLR3-LCD. (**a**) Cell death (PI positive cells) and lysosomal membrane permeabilization (acridine orange low) profile of SH-SY5Y cells expressing either wt MLKL or dn MLKL mutants (strep-MLKL and PsKD) following treatment with IFN/pIC for 24 h. Data points represent the mean □±□ S.E.M. of n = 4. ****p < 0.0001; ***p = 0.0008; **p= 0.0043. (**b**) Confocal images of SH-SY5Y cells expressing LAMP-1-RFP treated or not with IFN/pIC for 24 h stained with CellMask and Hoechst. Representative images from n = 3 are shown as merge (LAMP-1-RFP, CellMask green and Hoechst). Scale bars 10 µm. (**c**) Quantitative analysis of (**b**). Intracellular distribution profile of LAMP-1-RFP associated fluorescence intensity comparing the perinuclear with the peripheral area of SH-SY5Y cells expressing LAMP-1-RFP. Data points represent the mean □±□ S.E.M. At least 40 cells were analyzed per condition from n = 2. ****p < 0.0001. (**d**) Histogram overlays LAMP-1-Flag surface staining in DAPI negative subset of SH-SY5Y cells expressing LAMP-1-Flag. Left panel overlay 2^nd^ antibody only control with untreated; right panel overlay 2^nd^ antibody control with IFN/pIC treated for 24 h condition. Representatives of n = 3 are shown. (**e**) Quantitative analysis of (**d**). Fold increase of surface LAMP-1 associated fluorescence in DAPI negative subset of SH-SY5Y cells treated with IFN/pIC for the indicated times. Data points represent the mean □±□ S.E.M. of n = 3. *p = 0.043; **p = 0.0037; ***p = 0.0005. (**f**) Western Blot analysis of WCE and surface MLKL using anti-MLKL, -LAMP-1, -Na/K ATPase (Na/K), and -Strepdavidin (strep) antibodies. SH-SY5Y cells were treated for the indicated times with IFN/pIC and labeled with surface biotin or not followed by Streptactin pull down (IP biotin). WCE= whole cell extract. Representative experiment of n = 3 is shown. (**g**) Cell death profile of SH-SY5Y cell left untreated or treated with IFN/pIC in combination with MK6-83 for 48 h. Data points represent the mean □±□ S.E.M. of at least n = 4. * p = 0.0411; ****p < 0.0001. (**h**) Cell death profile of SH-SY5Y cell left untreated or treated with IFN/pIC in combination with Vacuolin-1 (VAC) or LY294002 (LY) for 48 h. Data points represent the mean □±□ S.E.M. of n = 4. ****p < 0.0001.

### ESCRT balances TLR3-LCD to maintain survival

ESCRT-dependent membrane repair can interfere with programmed cell death induction ^6,20^. Manumycin A, a farnesyltranserase inhibitor known to interfere with ESCRT activity increased IFN/pIC induced cell death and supported a pro-survival role of ESCRT during TLR3-LCD (Fig. 3a). To confirm these results, we over-expressed a component of the ESCRT-III complex, called charged multivesicular body protein (CHMP)3. While CHMP3^wt^ does not interfere with ESCRT-dependent membrane repair, a CHMP3 1-179aa fragment (CHMP3^mut^) can assemble the ESCRT-III protein complex at the side of membrane lesions, but fails to disassemble to finalize ILV formation ^21^. Since stable over-expression of CHMP3^mut^ is toxic at basal level ^20^, the expression was induced by doxycycline at the same time as pIC treatment (Suppl. Figure S5A). Expression of CHMP3^wt^ protected cells from IFN/pIC induced cell death, up to 48 h of treatment (Fig. 3b and c). In contrast, overexpression of CHMP3^mut^ sensitized cells to TLR3-LCD (Fig. 3c). Transferring the CHMP3 overexpression system to caspase-8 deficient HeLa cells, which undergo TLR3-LCD following IFN/pIC treatment, confirmed the protective role of ESCRT machinery during TLR3-LCD in an additional cell line (Suppl. Figure S5b-d). Thus, ESCRT balances TLR3-LCD induced cell death and maintains cell survival. The activity of the ESCRT machinery is visualized by the formation of a punctate pattern, which was obvious following IFN/pIC treatment ^22^ (Fig. 3d). Importantly, CHMP3^wt^ puncta overlapped with LAMP-1-RFP, which was used as a lysosomal marker. Likewise, CHMP3^mut^-GFP displayed a punctate pattern which co-localized with LAMP-1, albeit more pronounced than CHMP3^wt^ (Fig. 3d and e). Since CHMP3^mut^ lacks membrane repair function and fails to disassemble, the highly dynamic ESCRT machinery is sequestered at places of action. Thus, ESCRT-dependent membrane repair takes place at the lysosomes during TLR3-LCD, which is in line with MLKL-mediated LMP. In addition, ESCRT appeared to repair perinuclear lysosomes, since LAMP-1 as well as CHMP3^mut^ were more distributed to central region following IFN/pIC treatment (Fig. 3f; Suppl. Figure S6). However, despite the lysosomal clustering at the cell center, lysosomal MLKL was detectable at the plasma-membrane, when CHMP3^mut^ was overexpressed (Fig. 3g), emphasizing the pro-death role of lysosomal exocytosis by positioning lysosomal MLKL at the plasma-membrane.

**Fig. 3.**
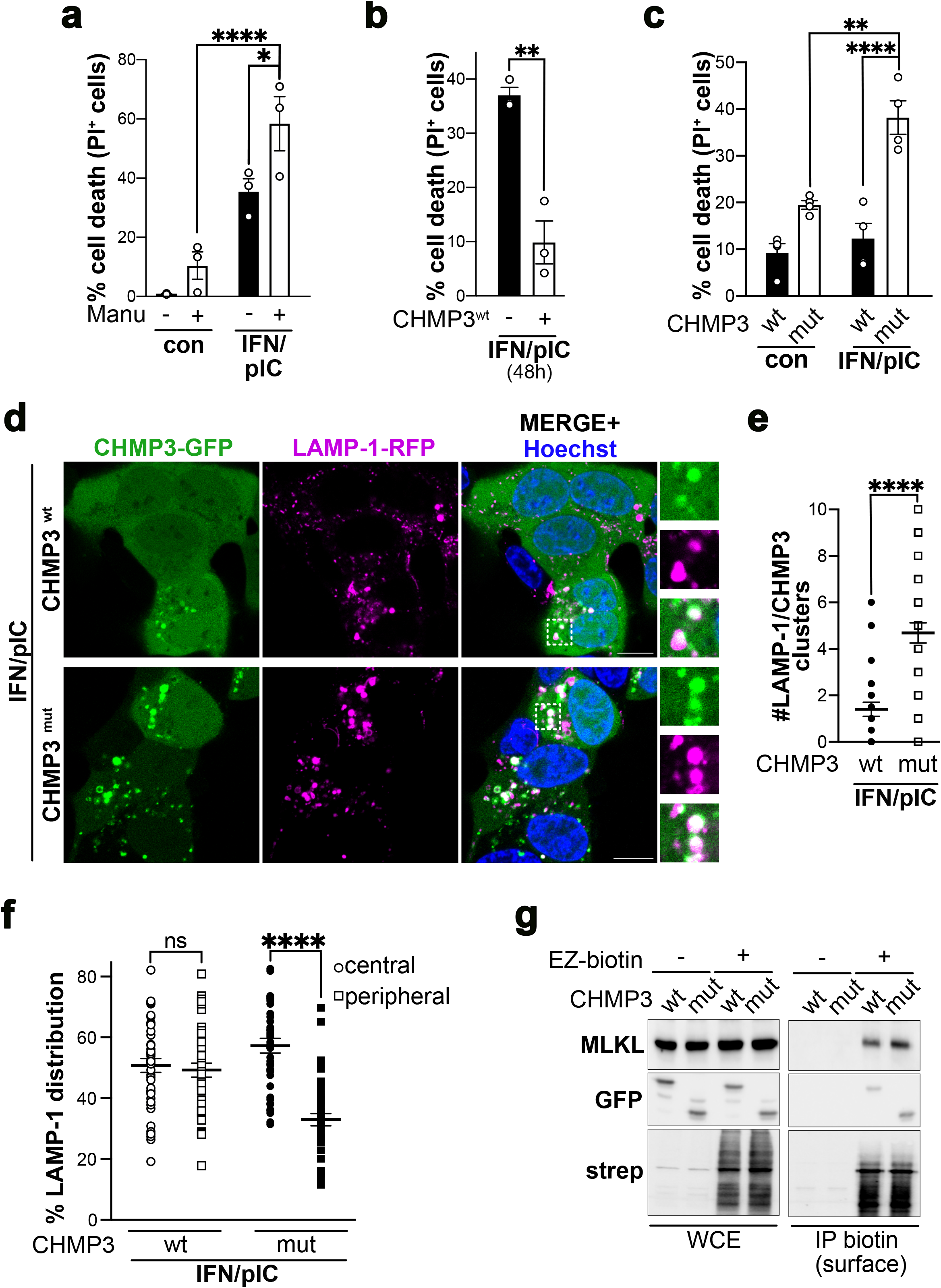
ESCRT balances TLR3-LCD to maintain survival. (**a**) Cell death profile of SH-SY5Y cells left untreated or treated with IFN/pIC for 48 h and co-treated with Manumycin A (Manu) by analyzing PI positive cells. Necroptosis was induced in HT-29 cells by treatment with pBZ for 12 h in combination with Manumycin A or not. Data points represent the mean □±□ S.E.M. of n = 3. ****p < 0.0001; *p = 0.0496; **p = 0.0013. (**b**) Cell death profile of CHMP3^wt^ overexpressing SH-SY5Y cells with or without Doxycycline following treatment with IFN/pIC for 48 h. Data points represent the mean □±□ S.E.M. of n = 3. **p =0.003. (**c**) Cell death profile of CHMP3^wt^ or CHMP3^mut^ expressing SH-SY5Y treated with IFN/pIC for 24 h analyzing PI positive cells. Data points represent the mean □±□ S.E.M. of n = 4. **p = 0.0016; ****p < 0.0001. (**d**) Confocal images of CHMP3^wt^- or CHMP3^mut^-GFP and LAMP-1-RFP expressing SH-SY5Y cells treated for 16 h with IFN/pIC and stained with Hoechst; merge of CHMP3-GFP, LAMP-1-RFP with Hoechst right panel). Inset is magnified. Scale bars 10 µm. Representative images of n = 2 are shown. (**e**) Quantification of confocal images in (**d**) counting the number of LAMP-1-RFP/ CHMP3^wt^-GFP or CHMP3^mut^-GFP clusters formed. Data points represent the mean □±□ S.E.M. of n = 2 analyzing at least 40 cells. ****p < 0.0001. (**f**) Intracellular distribution profile of LAMP-1-RFP associated fluorescence intensity in SH-SY5Y cells expressing LAMP-1-RFP and CHMP3^wt^-GFP or CHMP3^mut^-GFP treated for 16 h with IFN/pIC. Data points represent the mean □±□ S.E.M. At least 40 cells were analyzed per condition from n = 2. ns = not significant; ****p < 0.0001. (**g**) Western Blot analysis of WCE and surface MLKL using anti-MLKL, -GFP, and -Strepdavidin antibodies. SH-SY5Y cells overexpressing CHMP3^wt^-GFP or CHMP3^mut^-GFP were treated for the indicated times with IFN/pIC and labeled with surface biotin or not, followed by Strep-Tactin pull down. Representative experiment of n = 2 is shown.

### ESCRT integrates lysosomal MLKL into EVs

The ESCRT machinery is a key mediator of multivesicular bodies (MVB) biogenesis by promoting the formation of ILVs ^21^. Electron microscopy revealed that lysosomes did not only enlarge following IFN/pIC treatment as we previously observed, but were also charged with ILVs ^23^ (Fig. 4a). The average size of lysosomal ILVs was around 50 nm and thus comparable to the size of ILVs within MVB ^24^ (Suppl. Figure S7a). Fusion of MVB with the plasma-membrane releases ILVs the extracellular space as exosomes, which encompass a size average of 50– 150□nm ^25^. Accordingly, fusion of lysosomes with the plasma-membrane should release lysosomal ILVs in the extracellular space. Nanoparticle tracking analysis (NTA) of extracellular vesicles (EVs) revealed that IFN/pIC treatment increased the concentration of EVs secreted, without altering their size of around 100 nm (Fig. 4b and c; Suppl. Figure S7b). This is in sharp contrast to necroptotic plasma-membrane EVs, which encompass around 190□nm in size ^26^. Western blot analysis of EVs showed an increase in both MLKL as well as the lysosomal marker protein LAMP-1 following IFN/pIC treatment (Fig. 4b; Suppl. Figure S7c). Importantly, the classical exosome marker Flottilin-1 did not change following IFN/pIC treatment, reinforcing the idea that during TLR3-LCD not exosomes, but EVs with lysosomal origin are secreted. Thus, atypical activation of MLKL during TLR3-LCD results in the increased release of EVs containing lysosomal MLKL. To understand the relationship between EVs secreted following IFN/pIC treatment and lysosomal ESCRT activity, we used the previously established CHMP3 doxycycline-inducible overexpression system and analyzed EVs by NTA following sub-lethal IFN/pIC treatment for 16 h (Fig. 4e and f; Suppl. Figure S5a). SH-SY5Y cells expressing CHMP3^mut^ secreted significantly less EVs than CHMP3^wt^ expressing cells. Importantly, LAMP-1 as well as MLKL levels in EVs varied in correlation with EVs concentration, while Flottilin-1 levels remained unchanged comparing CHMP3^wt^ with CHMP3^mut^ expressing cells (Fig. 4g). In this way, MLKL as well as LAMP-1 were most prominent in EVs isolated from SH-SY5Y cells expressing CHMP3^wt^ and following treatment with IFN/pIC. Collectively, the ESCRT machinery packs lysosomal MLKL into lysosomal ILVs, which are then released into extracellular space as EVs to maintain survival. In contrast, in ESCRT-deficient cells MLKL accumulates at the lysosomes, then reaches the plasma-membrane via lysosomal exocytosis, where it ultimately executes its deadly function (Fig. 4h).

**Fig. 4.**
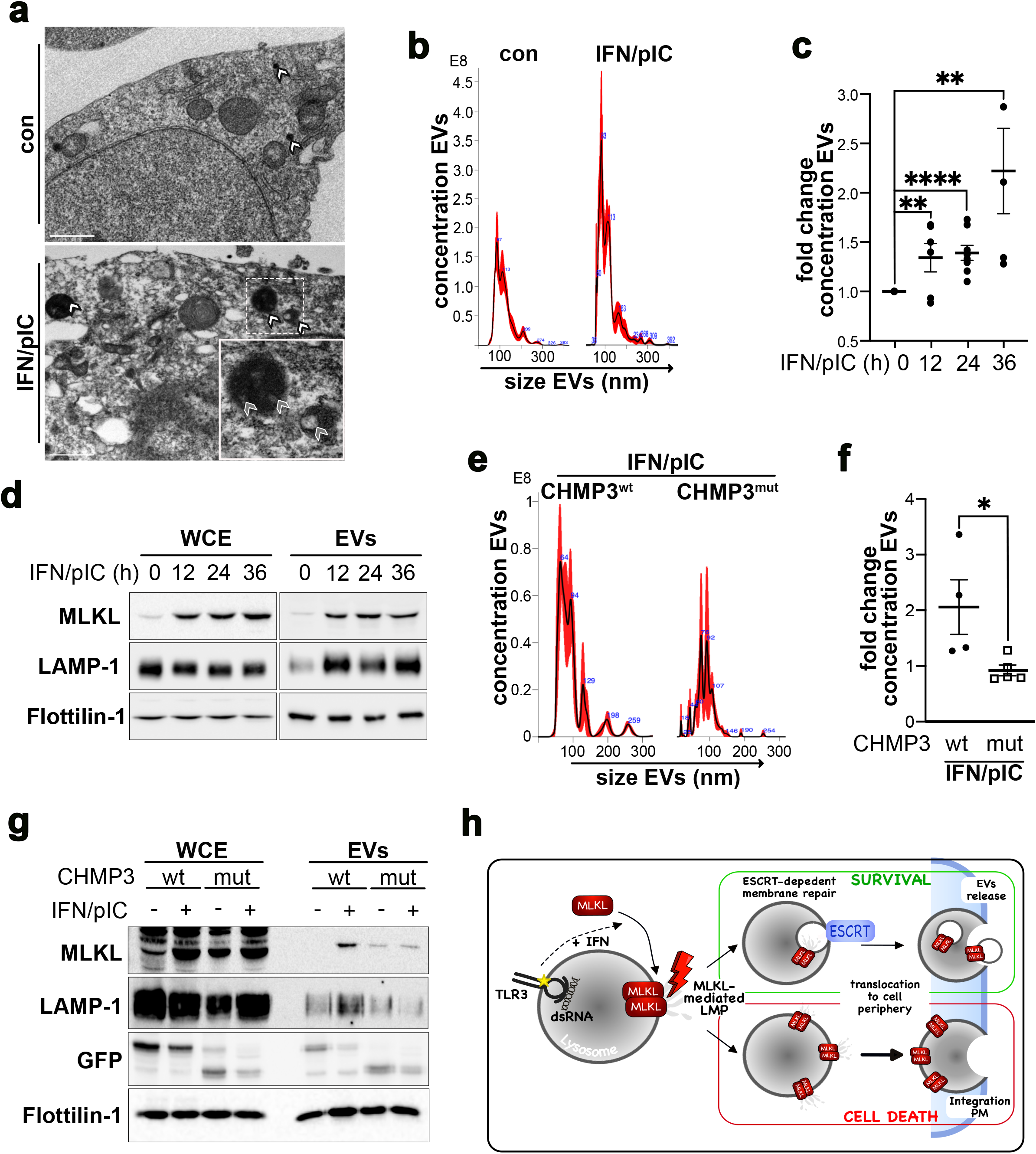
ESCRT integrates lysosomal MLKL into EVs. (**a**) Electron microscopic images of SH-SY5Y cells treated or not with IFN/pIC for 24 h. Insets are magnified. White arrowheads indicate lysosomes, grey arrowheads indicate luminal vesicles within lysosomes. (**b**) Size (nm)/ concentration (particles/ml) profile of extracellular vesicles (EVs) isolated from SH-SY5Y cells following treatment with IFN/pIC for 24h. A representative of n = 5 is shown. (**c**) Concentration profile of the fold increase of EVs isolated from SH-SY5Y cells treated with IFN/pIC for the indicated time. Data points represent the mean □±□ S.E.M. of at least n = 4. **p = 0.0071; 0,0011; ****p < 0,0001. (**d**) Western blot analysis of WCE (whole cell extract) and EVs isolated from SH-SY5Y cells treated for the indicated times with IFN/pIC using anti-MLKL, -LAMP-1 and Flottilin-1 antibodies. A representative of n = 3 is shown. (**e**) Size (nm)/ concentration (particles/ml) profile of extracellular vesicles (EVs) isolated from SH-SY5Y cells expressing CHMP3^wt^ or CHMP3^mut^ following treatment with IFN/pIC for 16h. A representative of n = 5 is shown. (**f**) Profile of fold change in concentration of EVs isolated from SH-SY5Y cells expressing CHMP3^wt^ or CHMP3^mut^ treated with IFN/pIC for 16 h. Data points represent the mean □±□ S.E.M. of at least n = 5. *p = 0.0381. (**g**) Western blot analysis of 10% WCE and EVs isolated from SH-SY5Y cells expressing CHMP3^wt^ or CHMP3^mut^ treated with IFN/pIC for 16 h using anti-MLKL, -LAMP-1 -GFP (CHMP3) and -Flottilin-1 antibodies. A representative of n = 2 is shown. (**h**) Schematic presentation of MLKL-induced TLR3-LCD and its modulation by ESCRT.

## Discussion

Maintaining the integrity of lysosomes is essential for cellular homeostasis, while LMP is associated with degenerative diseases, infection and cancer ^27^. Considering that LMP is a progressive process initiated by limited membrane permeabilization, cells have established safe-guard mechanisms to prevent deadly lysosomal membrane rupture ^28^. In this way, the ESCRT machinery can detect and repair small proton-permeable ruptures in the lysosomal membrane, to maintain lysosomal integrity and survival ^7,8,29^. The activity of the ESCRT machinery commonly covers membrane budding and scission, influencing next to membrane repair an enormous variety of cellular processes ^30^. Accordingly, we could observe ILVs formed within the lumen of lysosomes, which are subsequently released as EVs in an ESCRT-dependent manner.

A factor that will determine the level of LMP and the possibility for the ESCRT-machinery to repair membrane ruptures, will be the size of the pore inducing LMP. Our results show that LMP following IFN/pIC treatment is dependent on the executer of necroptosis, MLKL. Despite its central role in necroptosis, the mechanism by which MLKL induces plasma-membrane permeabilization remains unclear. However, several studies support the notion that MLKL can form cation channels in the plasma-membrane, which are responsible for membrane disruption and necroptosis ^31,32^. In analogy, translocation of MLKL to other cellular organelle membranes, such as the lysosomes, and subsequent pore formation within the lysosomal membrane could explain the observed MLKL-mediated LMP. From a regulatory point of view, translocation to different membrane compartments within a cell could be achieved by different MLKL post-translational modifications ^33^. In agreement, we show that during TLR3-LCD MLKL is phosphorylated in an atypical, RIPK3-independent manner on sites distinct from the classical necroptotic phospho-sites. By providing a potential therapeutical target to modulate cell fate outcomes in disease settings which arise from a disturbed life-death balance, it will be of great interest to identify the kinase phosphorylating MLKL during TLR3-LCD. In any case, the damages that MLKL causes at the lysosomal membrane appears to be limited, since the ESCRT machinery can repair them and maintain survival. Thus, MLKL presumably also only forms cation channels to mediate limited LMP and not pores large enough to release lethal lysosomal content, such as cathepsins into the cytosol.

The question arises of what is the trigger converting limited MLKL-mediated lysosomal ruptures into TLR3-LCD. One hypothesis could be that over the course of treatment the concentration of lysosomal MLKL increase. This might not only be true for one lysosome per se, but for the total number of lysosomes affected. In any case, the membrane repair capacities of the ESCRT machinery will saturate and be overwhelmed, not able to ensure efficient lysosomal membrane repair anymore. Exceeding the MLKL threshold at the lysosomes could be mediated by the combination of IFN-I and pIC treatment, while additional TLR3-mediated IFN production could contribute to cell death induction ^34,35^. Damaged and unrepaired lysosomes undergo increased lysosomal exocytosis and likewise integrate the lysosomal MLKL-pore into the plasma-membrane. In analogy to necroptosis, this would result in osmotic imbalance, which goes along with the cellular swelling, that we observed during TLR3-LCD. Considering that plasma-membrane rupture is an active process ^36^, the lysosomal MLKL-pore might also actively be converted into a membrane disrupting pore at the plasma-membrane to execute TLR3-LCD. The precise mechanism how active lysosomal MLKL, however, triggers an enhanced mobility of lysosomes to the cell periphery to execute cell death will have to be elucidated in further studies.

Last, this study also highlights that MLKL-mediated membrane permeabilization is not a point of no return in death signaling, but instead can provide a survival signal. As an interferon-sensitive gene, MLKL might fulfill a still unknown anti-viral function in order to protect the host from viral infection. Accordingly, MLKL-mediated LMP recruits and activates the ESCRT machinery, which is commonly associated with viral infections. Recognition and sensing of viral infection by MLKL could be mediated by direct binding of MLKL ^37^. In this context, it is tempting to speculate that EVs loaded with lysosomal MLKL could provide a danger signal and communicate viral infection within the cellular microenvironment. However, further experiments will be required to pinpoint the pro-survival, anti-viral role of MLKL.

## Materials and Methods

### Cell culture and treatment procedure

SH-SY5Y, HT-29 and HEK293T cells (all purchased from ATCC) were cultured in Dulbecco’s modified eagle’s minimal essential medium. Cell cultures were routinely tested for mycoplasma contamination. All cells were cultured in low glucose DMEM or McCOY’s medium (Sigma, Germany) supplemented with 10% FBS (ThermoFisher, Germany), and grown in a humidified incubator containing 5% CO_2_ at 37□°C. The cells were frequently passaged at sub-confluence, and seeded at a density of 0.3–0.5□×□10^6^ cells/ml. To induce TLR3-LCD, SH-SY5Y cells were pre-treated for 16 h with Type I Interferon (1000 Units/mL; R&D Systems, 11200-2) and treated with HMW Poly(I:C) (10 μg/mL; Invivogen, tlrl-pic). To induce necroptosis HT-29 cells were pretreated for 1 h with BV6 (2,5 μM; Cliniscience, HY-16701) and 10 μM z-Val-Ala-DL-Asp(Ome)-fluoromethylketone (zVAD-fmk; Cliniscience, A12373) followed by treatment with Poly(I:C) for 12h. Other inhibitors included RIPK3 inhibitor GSK872’ (5 μM; Selleckchem; S8465), Necrosulfonamide (NSA; 5 μM; Selleckchem, SE-S8251), MK6-83 (10 μM; Sigma, SML-1509); LY294002 (5 μM; Cliniscience; HY-10108); Vacuolin-1 (5 μM; Calbiochem, CAS 351986-85-1); Manumycin A (2,5 μM; Enzo, ALX-350-241-M001); BAPTA-AM (5 μM; Selleckchem; SE-S7534);

The lentiviral CHMP3^wt^ and ^mut^ constructs were kindly provided by Prof. Petr Broz, University Lausanne, Switzerland; the lentiviral MLKL-GFP and dn MLKL-GFP constructs by Prof. Peter Vandenabeele, VIB Ghent, Belgium. To generate stably expressing SH-SY5Y cells, viruses were produced in HEK 293T cells by transfecting the corresponding plasmid o interest as well as with pVSVg (Addgene, #8454) and psPAX2 (Addgene, #12260) using Lipofectamine 2000 (Thermo Fisher Scientific, 11668019) according to the manufacturer’s instructions. Twenty-four and 48 h later, virus-containing supernatant was collected, filtered and used to infect SH-SY5Y cells. Two days later, the transduced cells were selected by growth in the appropriate antibiotic. For Crispr Cas9 mediated knock out, the oligos containing the gene-specific sgRNA target were cloned into the LentiCRISPRv2 Blasticidin (Addgene, #83480). The CRISPR/Cas9 primer sequences were as followed:

guide 1: 5’-GAACCGCTTCAAGGCTGCCC *TGG*-3’;

guide 2; 5’-CACACCGTTTGTGGATGACC *TG*-3’.

The following primary antibodies were used for western blotting: MLKL (CST; 14993S); p-MLKL (CST; 916895); LAMP-1 (CST; 9091S); Flottilin-1 (BD; 610820); FLAG (Sigma-Aldrich; F1804); GFP (Abcam; ab6673); Na/K ATPase (CST; D4Y7E); Strepdavidin-HRP (Sigma-Aldrich; OR03L) and actin-HRP (Sigma-Aldrich, A3854).

### Cell death and lysosomal permeabilization assay

Cells were stained with 5 μM Propidiumiodide (PI; Life Technologies) and cell death measured by flow cytometry analyzing PI positive population of floating and adherent cells in the FL-2 channel. Flow-cytometric analyses were conducted using the FACSCalibur and data analyzed using the Flowjo software. Alternatively, cell death/PI positivity was analyzed with an IncuCyte ZOOM system (Sartorius). Percent cell death was calculated as followed:

[100 × (inducedfluorescence–backgroundfluorescence)] ÷ (maximalfluorescence achieved by Triton-X-100 permeabilization – background fluorescence). Lysosomal stability was assayed using acridine orange (AO) (Sigma-Aldrich, Saint-Louis, MO, USA) relocation methods. SH-SY5Y cells were loaded with AO (10 μg/mL) for 15 minutes in complete culture medium at 37 °C prior the end of the treatment. The increase in green fluorescence due to the release of AO from permeabilized lysosomes or the decrease of AO-associated red fluorescence was quantified by flow cytometry in the FL1 and FL2 channel, respectively. The data are presented as mean □±□standard error of the mean (S.E.M) of at least three independent experiments. For other statistical analysis, two-way or one-way ANOVA; or two-tailed unpaired *T*-test was used. \**P*□<□0.05 was considered as significant.

### MLKL activation

For detection of oligomeric MLKL cells were directly lysed in 2x sample buffer and sonicated with a needle sonicator (10 sec, 40% amplitude). Samples were run under nonreducing (w/o DTT) and reducing conditions (+ 0,1 M DTT) on SDS-PAGE. Phosphorylation was determined by running TCA precipitated lysates on Phostag SDS-PAGE (FUJIFILM, AAL-107) according to manufactur’s protocol. For lambda phosphatase treatment cells were lysed in RIPA buffer and incubated with lambda phosphatase (New England biolabs; P0753S) for 30 min at 37 °C prior to TCA precipitation and loading on Phostag SDS-Page. 400U λ PPase were added prior to TCA precipitation when indicated. Enzymatic reactions were allowed to proceed for 30□min at 37□°C and subsequently.

### Lysosomal positioning and exocytosis

SH-SY5Y cells stably overexpressing LAMP-1-Flag-RFP (Addgene; #102931) and stained with CellMask Green (1/1000; Invitrogen, C37608). Cells were imaged by confocal microscopy using a LSM880 confocal microscope (Zeiss) and images (8□bit) were acquired with a Plan-Apochromat 63X/1.4 oil objective at a resolution of 1528 by 1528 pixels (pixel size 70 nm□×□70 nm). Images were processed blinded using Fiji imaging software and intracellular distribution was determined as previously described (*36*). Briefly, nuclear size was measured and perinuclear area determined by multiplying nuclear diameters by 1,5. Fluorescence intensity of LAMP-1-FLAG-RFP within the perinuclear area was measured and subtracted from the total LAMP-1 associated fluorescence intensity to obtain the peripheral lysosomal distribution. For each condition, 7–9 fields of view were randomly selected and imaged from at least two independent experiments. Cells within these fields were analyzed, resulting in 5–10 data points per condition. All quantitative datasets in this publication are presented as averages ± S.E.M. For lysosomal exocytosis SH-SY5Y cells stably overexpressing LAMP-1-Flag-RFP were treated with IFN/pIC and only adherent cells collected. Surface LAMP-1-Flag-RFP was stained using anti-Flag antibody at 4 °C for 4 h washed in FACS buffer (PBS + 0.5 % BSA) and fixed with 2% PFA, before incubation with corresponding secondary Alexa 488 or Alexa 564 coupled antibodies (Life science; A21151 or A31571). LAMP-1-Flag-RFP surface staining was determined in the DAPI negative cell population by flow cytometry.

### Biotinylation of surface proteins

Adherent SH-SY5Y cells (25 x 10^6^ cells) were washed in ice cold PBS and incubated with membrane-impermeable biotinylation reagent, EZ-Link™ Sulpho-NHS-Biotin (1 mg/ml in PBS buffer, pH 8; Thermo fisher; 21217) was added (or not; control) and incubated for 2 h on ice with gentle shaking. The reaction was quenched by 100mM Glycine in PBS buffer and cells were lysed in RIPA buffer and sonicated. Precleared lysates were incubated with MagStrep “type3” Strep-Tactin beads (IBA; 2-1613-002) over night and 2,5 % WCE retained for analysis of the input fraction. The biotinylated proteins were eluted from the beads using SDS–PAGE sample buffer and separated on SDS–PAGE for immunoblot analysis.

### Extracellular vesicles

Supernatants of 10 x 10^6^ SH-SY5Y cells were collected and centrifuged at 2000 x g for 10 min to pellet cells. Supernatants were further centrifuged at 10 000 x g for 30 min to remove microvesicles. Finally, supernatants were centrifuged at 100 000 x g for 70 min in ultracentrifugation using type 70 Ti rotor (Oprima XL-80K, Beckman, Brea, CA, USA) to pellet the EVs. Obtained pellets were resuspended in 2 x sample buffer (0.1□M DTT, 60 mM Tris/HCl (pH 6.8) 2 % (w/v) sodium dodecyl sulphate (SDS), 15 % (v/v) glycerol, 0.05 % (w/v) Bromophenol Blue) and heated for 5 min at 95 °C prior to SDS-PAGE. For a single-particle concentration and size measurements, EV pellets were resuspended in PBS and analyzed using Nanoparticle Tracing Analysis (NTA) (NanoSight NS300, version 3.1.54, Malvern Instruments).

### Electron microscopy

For transmission electron microscopy SH-SY5Y were fixed with 2% glutaraldehyde (EMS) in 0.1 M sodium cacodylate (pH 7.4) buffer at room temperature for 30 min. After washing three times in 0.2 M sodium cacodylate buffer, cell cultures were post-fixed with 2% osmium tetoxide (EMS) at room temperature for 1 h and dehydrated in a graded series of ethanol at room temperature and embedded in Epon. After polymerization, ultrathin sections (100 nm) were cut on a UCT (Leica) ultramicrotome and collected on 200 mesh grids. Sections were stained with uranyl acetate and lead citrate before observations on a Jeol 1400JEM (Tokyo, Japan) transmission electron microscope, equipped with an Orius 600 camera and Digital Micrograph Software (Gatan).

## Acknowledgments

We thank M. Bellina, C. Vanbelle, M.S. Meszaros, E. Errazuriz-Cerda for technical advice and support, and S. Lebecque, L. Wong, B. Crocker, U. Ros and P. Broz for discussions and critical reading of the manuscript.

This work was supported by Centre Léon Berard ; La Ligue Contre le Cancer ; Institute Convergence PLAsCAN (ANR-17-CONV-0002) ; LabEX DEVweCAN (University of Lyon) ; Agence Nationale de la Recherche (ANR-18-CE13-0005-01) ; Fondation LEEM.

## Author contributions

Conceptualization: MC, GI, KW; Methodology: KW, JC, CG; Investigation: KW, JC, CG; Visualization: KW; Funding acquisition: MC, KW, GI; Project administration: CJ, MC, GI, KW; Supervision: KW, GI; Writing – original draft: KW; Writing – review & editing: MC, GI, KW

## Data and materials availability

All data are available in the main text or the supplementary materials.

