## Supplemental Figures for "Lysosomal MLKL is balanced by ESCRT to control cell death"

### Suppl. Figure S1

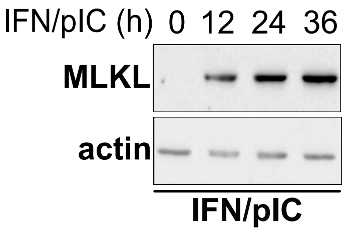


Western blot analysis of SH-SY5Y cells primed for 16 h with IFN-I followed by treatment with pIC for the indicated times using anti-MLKL and -actin antibodies.

#### Suppl. Figure S2

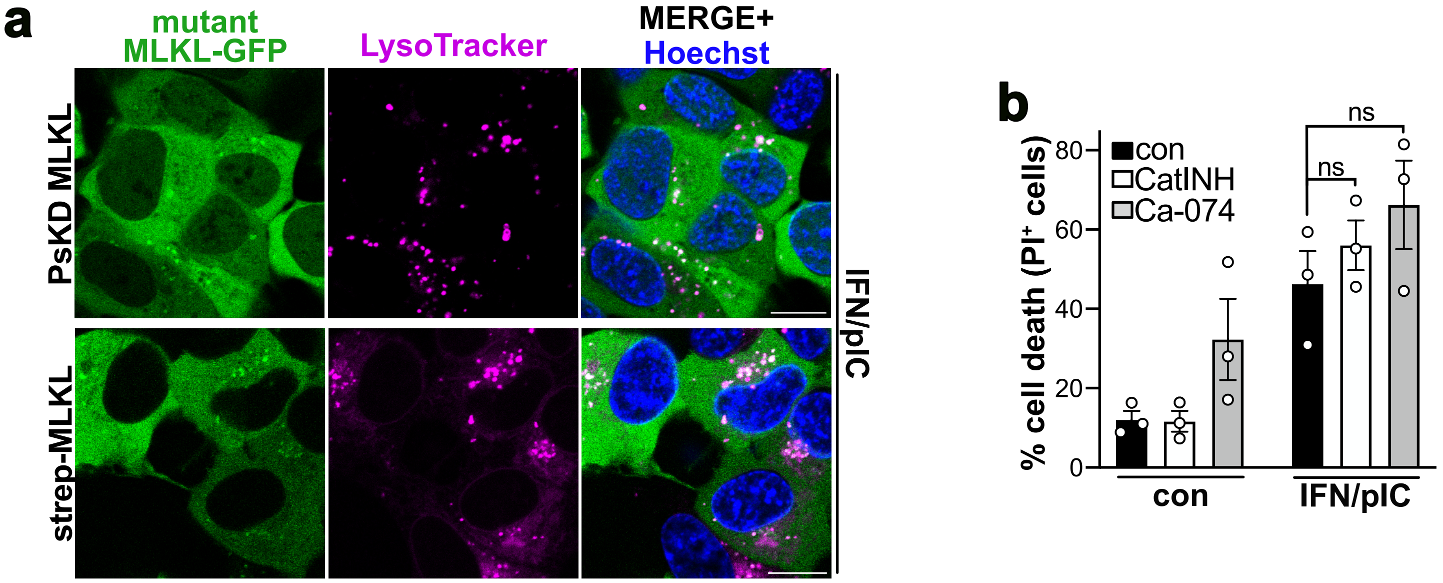


(**a**) Confocal images of SH-SY5Y cells expressing wt MLKL or the dominant negative (dn) MLKL mutants: Pseudokinase domain of MLKL (PsKD) and N-terminally strep-tagged MLKL, stained with LysoTrackerRed and Hoechst. Merged image of MLKL-GFP, Lysotracker and Hoechst right panel. Scale bars 10 µm. Representative images of n = 1 are shown. (**b**) Cell death profile analyzing the percentage of Propidium iodid positive (PI+) cells following treatment with IFN/pIC for 48 h in combination with the Cathepsin inhibitors CatINH, or Ca-074.

#### Suppl. Figure S3

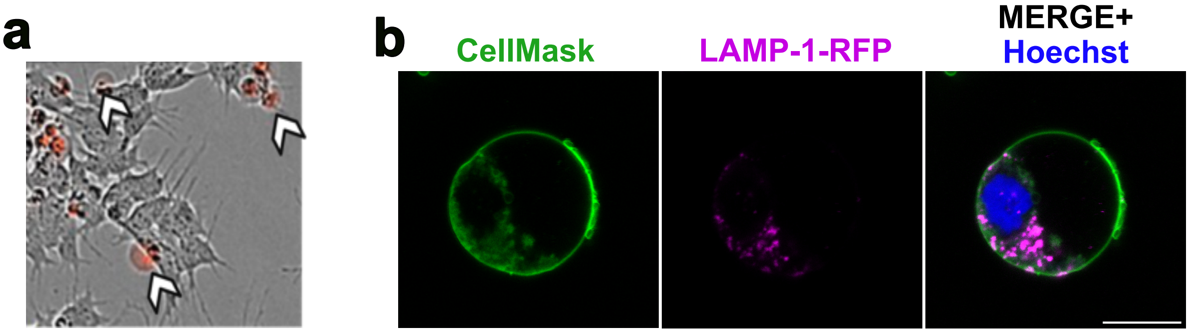


(**a**) Phase image merged with PI images of SH-SY5Y treated with IFN/pIC taken with Incucyte device. Balloon-like structures are indicated by arrow heads. (**b**) Confocal images of SH-SY5Y cells stably expressing LAMP-1-RFP and stained with CellMask treated for 24 h with IFN/pIC. Merge image with Hoechst is shown. A representative of n = 2 is shown. Scale bar 10 µm.

### Suppl. Figure S4

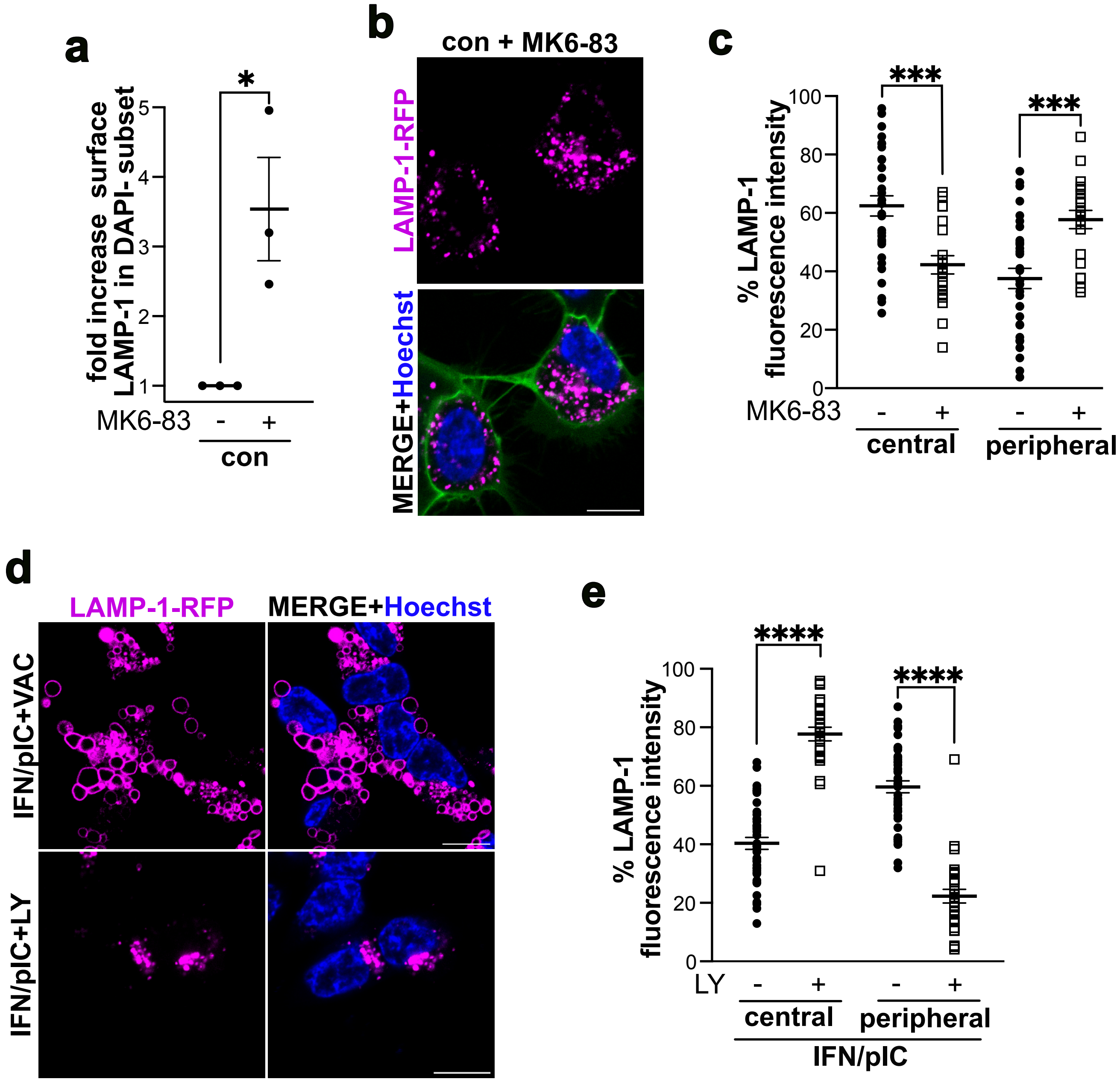


(**a**) Fold increase of surface LAMP-1 associated fluorescence in DAPI negative subset of SH-SY5Y cells treated with MK6-83 for 24 h. Data points represent the mean ± S.E.M. of n = 3. *p = 0.00266. (**b**) Confocal images of LAMP-1-RFP expressing SH-SY5Y cells treated with MK6-83 for 24 h and stained with CellMask green and Hoechst. Bottom panel merge of Hoechst, CellMask and LAMP-1-RFP. A representative of n = 2 is shown. Scale bar 10 µm. (**c**) Quantification of confocal images in (**b**). Profile of the intracellular distribution comparing the percentage of LAMP-1-RFP associated fluorescence intensity of the perinuclear and peripheral cellular area. Data points represent the mean ± S.E.M. of n = 2. Per condition at least 25 cells were analyzed. ***p = 0.0005. (**d**) Confocal images of LAMP-1-RFP expressing SH-SY5Y cells treated on the top panel with IFN/pIC and Vacuolin-1 and on the bottom panel with IFN/pIC and LY294002 both for 36 h and stained with Hoechst. Right panel merge of LAMP-1-RFP with Hoechst. Representative images of n = 2 are shown. Scale bars 10 µm. (**e**) Quantification of confocal images in (**d**) bottom panel: treatment with pIC/IFN and LY294002 for 36 h. Profile of the intracellular distribution comparing the percentage of LAMP-1-RFP associated fluorescence intensity of the perinuclear and peripheral cellular area. Data points represent the mean ± S.E.M. of n = 2. Per condition at least 30 cells were analyzed. ****p = 0.0001.

### Suppl. Figure S5

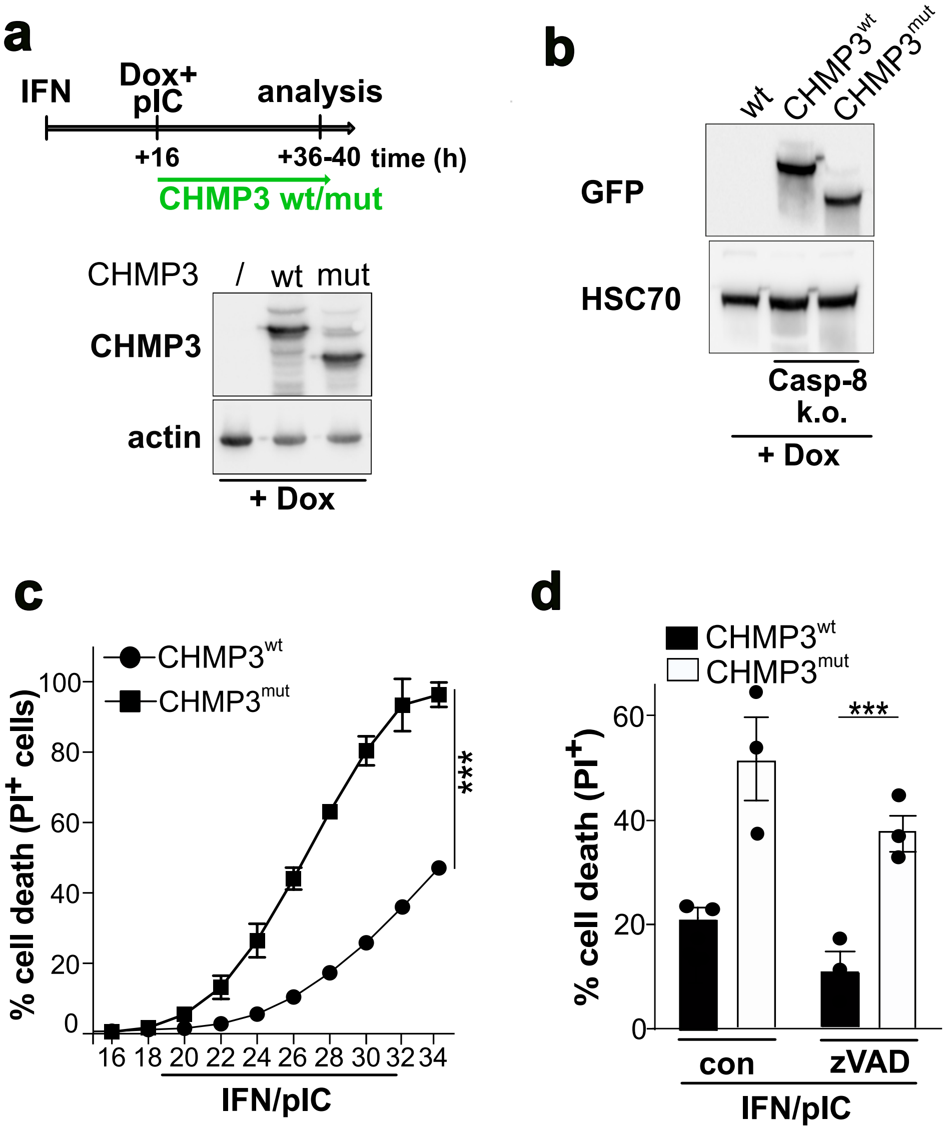


(**a**) Top: Schematic representation of time line for treatment of doxycycline (Dox) inducible CHMP3wt or CHMP3mut (dominant negative mutant spanning aa 1-179) experiments. Bottom: Western blot analysis of SH-SY5Y cells stably expressing control plasmid or GFP- CHMP3$^{wt}$ or GFP-CHMP3$^{mut}$ following treatment with 1µg/ml Dox using anti-GFP and -actin antibodies. A representative of n = 3 is shown. (**b**) Western blot analysis of CRISPR/Cas9 knock-out (KO) Caspase-8 HeLa cells stably expressing dox-induced CHMP3wtand mutGFP fusion proteins using anti-GFP and -HSC70 antibodies. (**c** and **d**) Cell death profile of Caspase-8 KO HeLa cells analyzing the percentage of PI+ cells following dox-induction and (**c**) IFN/pIC treatment for the indicated times or (**d**) following IFN/pIC treatment for 26 h in combination or not with 25 µM zVAD.

#### Suppl. Figure S6

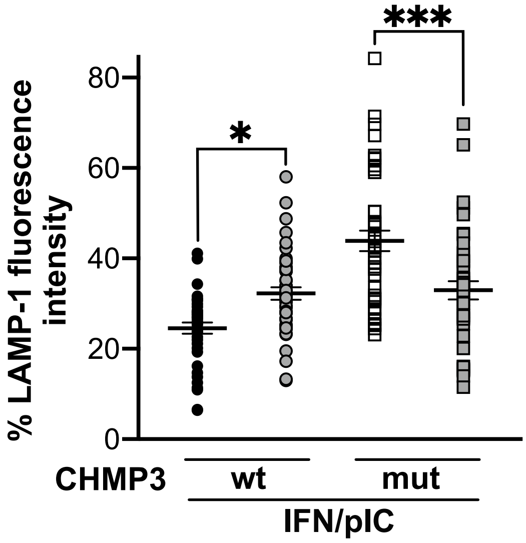


Intracellular distribution profile of CHMP3wtGFP or CHMP3[mut]-GFP associated fluorescence intensity in SH-SY5Y cells expressing LAMP-1-RFP and CHMP3[wt]-GFP or CHMP3[mut]-GFP treated for 16 h with IFN/pIC. Data points represent the mean ± S.E.M. At least 40 cells were analyzed per condition from n = 2. *p=0.014; ***p = 0.0001.

**Suppl. Figure S7**

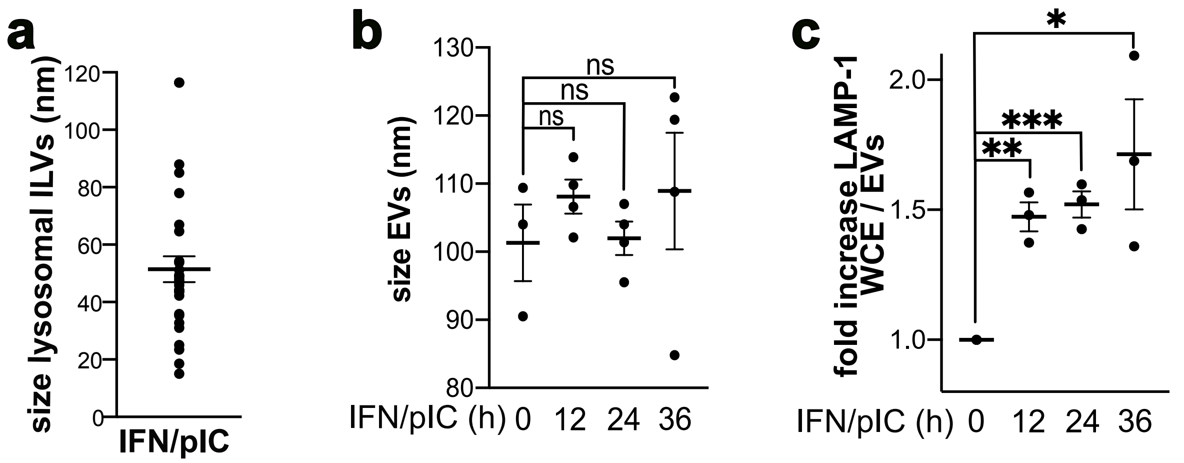


(**a**) Profile of size (diameter; nm) of lysosomal ILVs. (**b**) Profile of fold change in concentration of EVs isolated from SH-SY5Y cells treated with IFN/pIC for the indicated times. Data points represent the mean ± S.E.M. of n = 5. ns= not significant. (**c**) Profile of fold increase of densitometric analysis of LAMP-1 signal in WCE compared to EVs. Data points represent the mean ± S.E.M. of n = 3. *p = 0.028; **p = 0.0011; ***p = 0.0005.

#### **Suppl. Figure S8** - uncropped Western blots Fig 1

**a**

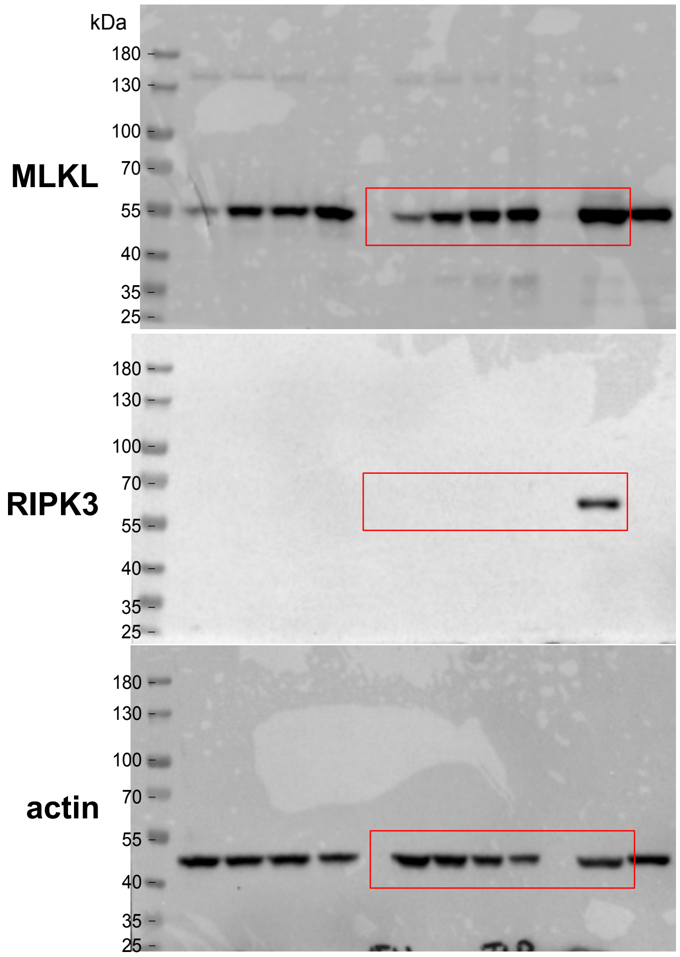


**b**

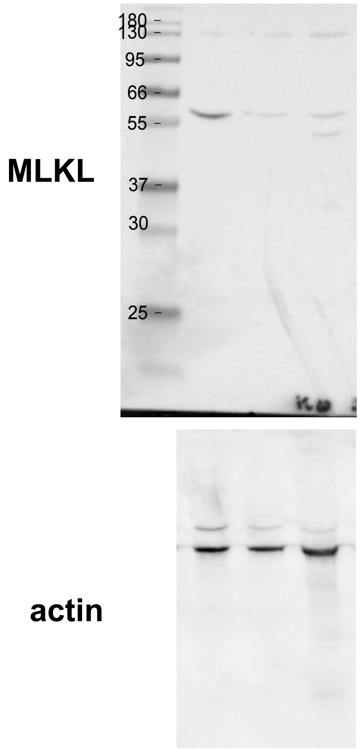


**e**

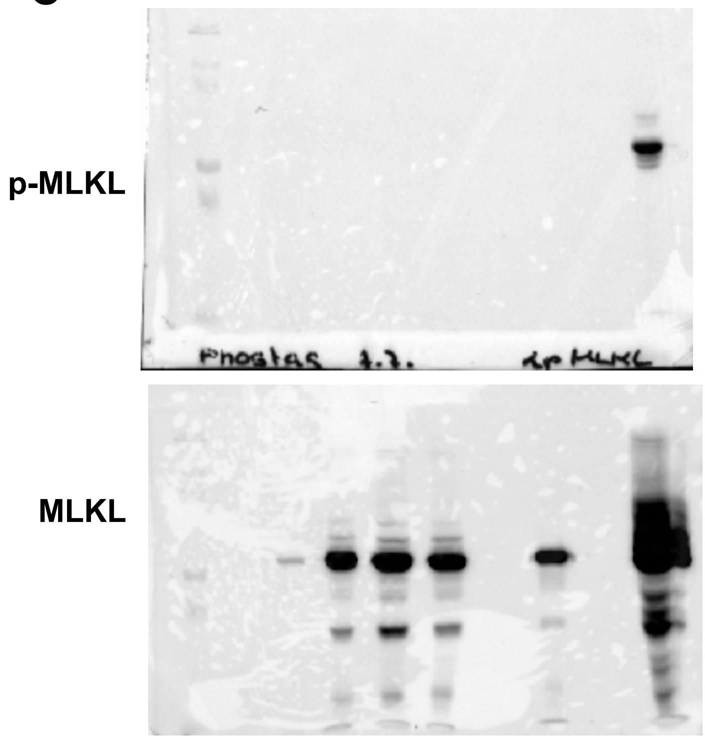


**f**

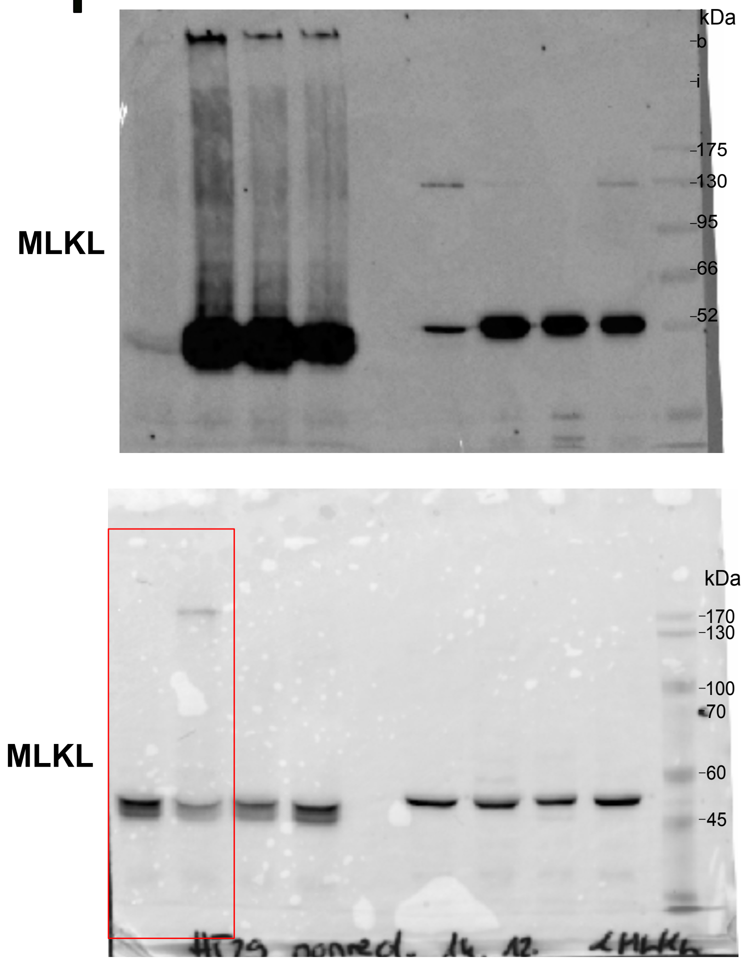

**Suppl. Figure S9** - uncropped Western blots Fig 2

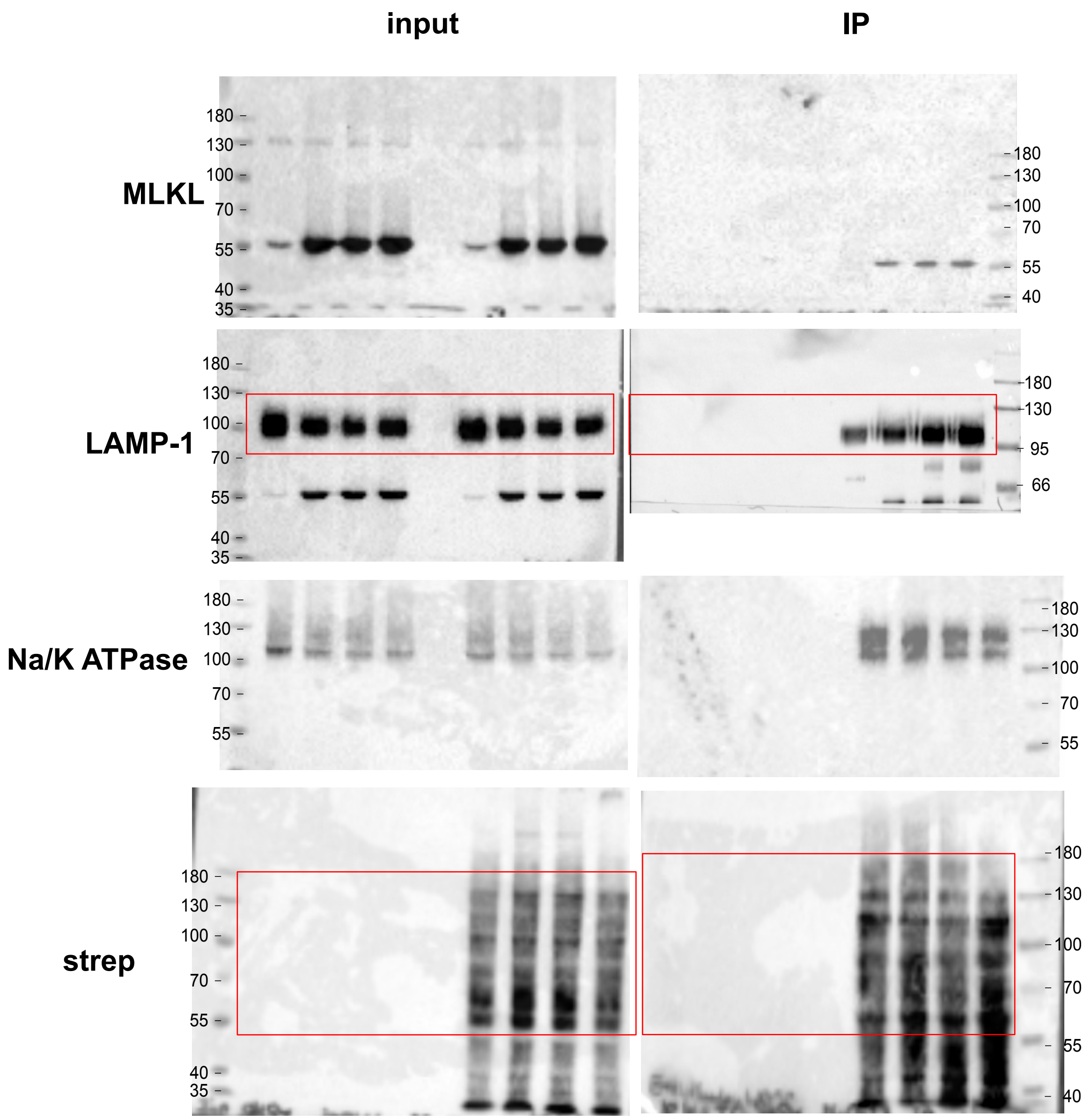

**Suppl. Figure S10** - uncropped Western blots Fig. 3

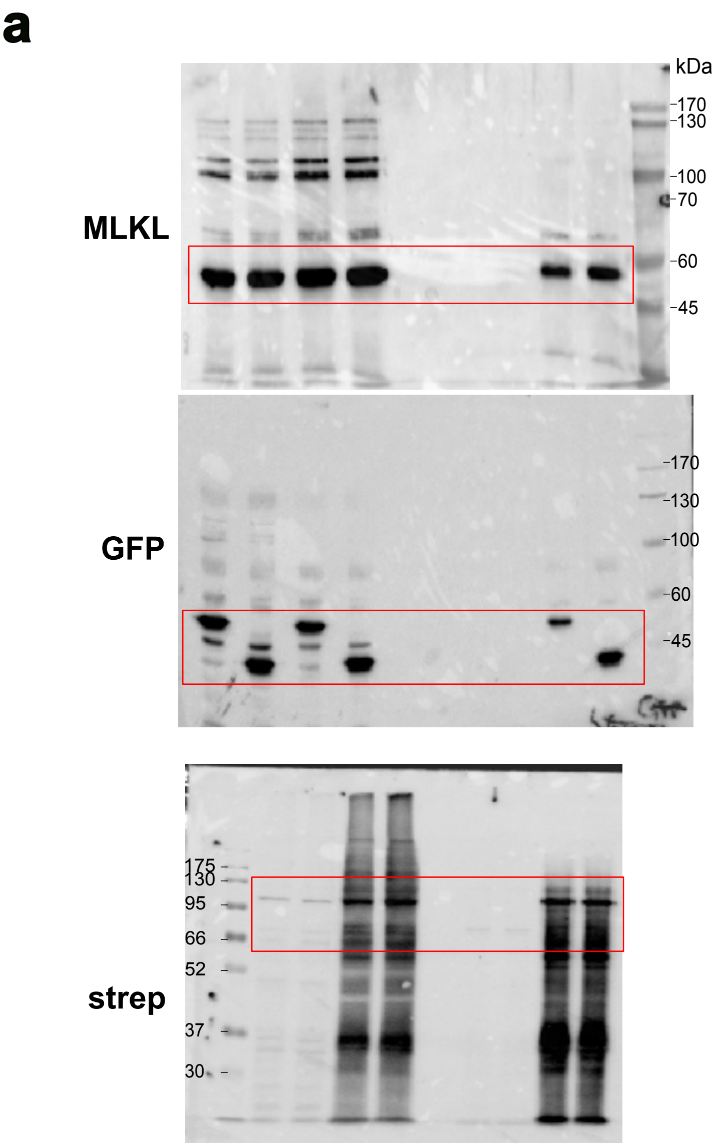

**Suppl. Figure S11** - uncropped Western blots Fig. 4

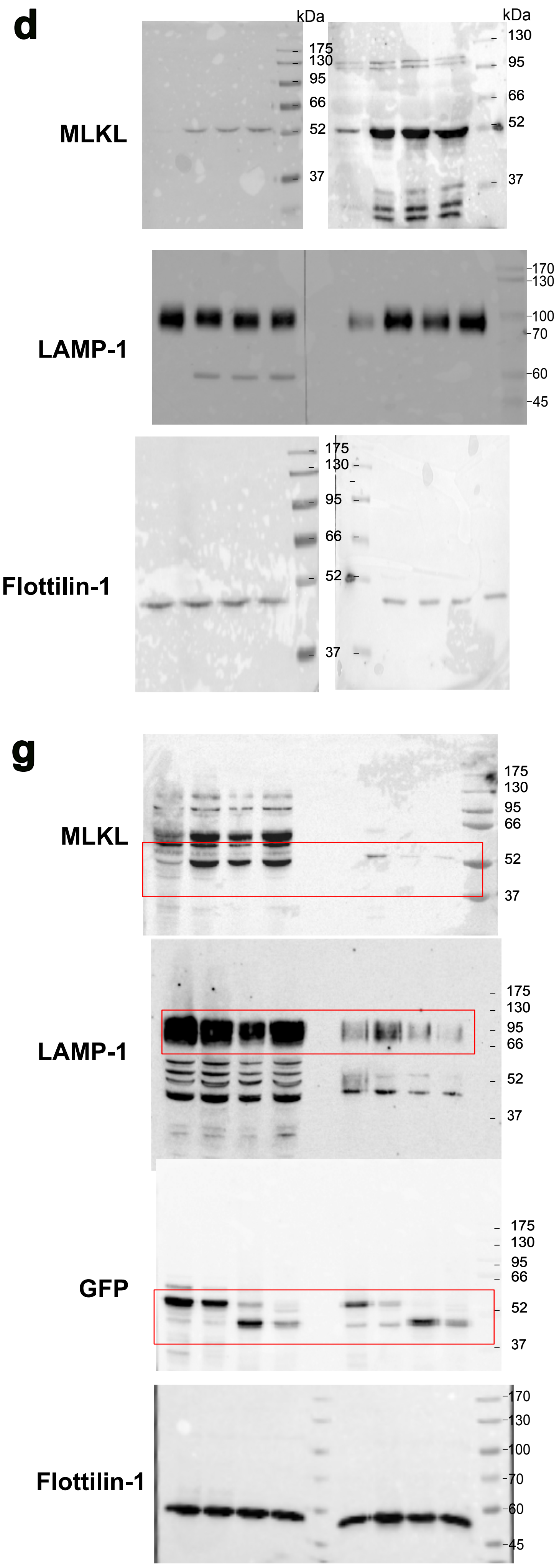

**Suppl. Figure S12** uncropped Western blots Suppl. Figures

#### Fig S1

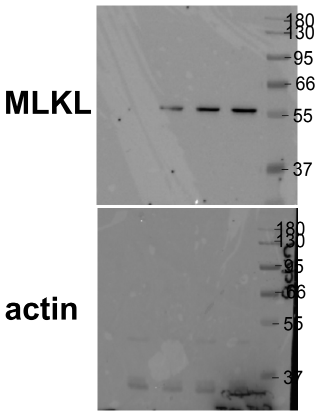


#### Fig S5 a

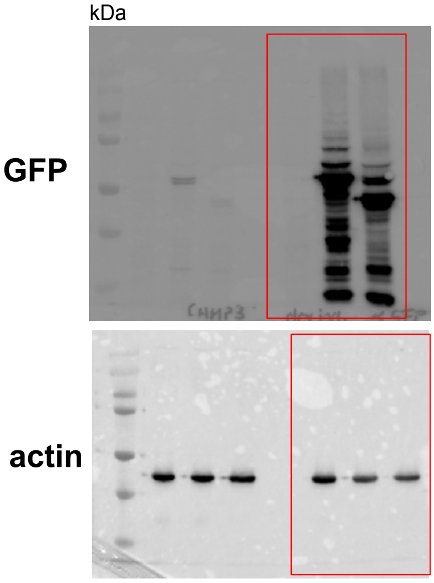


#### Fig S5 b

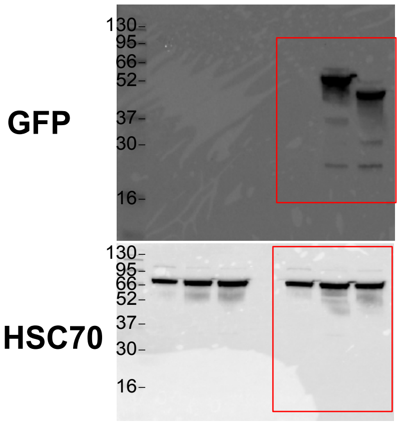
